# Population structure and aquatic environment shape kin discrimination in toad tadpoles

**DOI:** 10.1101/2025.08.07.669028

**Authors:** Kazuko Hase

## Abstract

Kin discrimination may vary with population genetic structure and environmental context. Many animals use olfactory cues in kin recognition, and learned recognition cues may be associated with major histocompatibility complex (MHC) genes. Genetic variation differs among lineages and local populations, and admixture may increase genetic variation within a population. Here, I compared kin discrimination in tadpoles from two local populations of the Japanese eastern common toad (*Bufo formosus*): a homogeneous population belonging to the eastern lineage and an admixed population containing both eastern and western lineages. Genetic analyses revealed higher MHC polymorphism in parental toads from the admixed population. Tadpoles were reared under four conditions combining social experience (group or solitary rearing) and water type (tap or pond water), and tested using binary-choice assays. Kin discrimination was detected only in group-reared tadpoles from the admixed population maintained in tap water, whereas no kin discrimination was detected in the homogeneous population under any tested condition. In a separate test of familiarity-based association, tap-water-reared tadpoles from the admixed population preferred familiar siblings when unfamiliar siblings had been reared in pond water. These findings highlight the importance of considering both population genetic structure and rearing context when investigating social recognition.

## 1. INTRODUCTION

Kin discrimination, the differential treatment of kin and non-kin, varies among species and can depend on developmental experience and environmental context [1–5]. In animals that use learned cues to identify relatives, discrimination may rely on phenotype matching, whereby individuals compare conspecifics with a recognition template acquired through previous social experience [6,7]. Unlike familiarity-based recognition, phenotype matching allows individuals to discriminate between unfamiliar kin and non-kin [8]. In vertebrates, olfactory cues are often involved in social recognition, and major histocompatibility complex (MHC) genes have been implicated in kin-biased association preferences [9–11]. However, the expression of kin discrimination may depend not only on recognition mechanisms but also on population genetic structure, and ecological conditions [2,12].

Anuran tadpoles provide a useful system for investigating these relationships. Tadpoles often show social aggregation for foraging efficiency [13,14], heat retention [15], and predator avoidance [16]. Many species exhibit kin-biased association through phenotype matching or familiarity-based recognition [3,4,17]. However, kin discrimination is not consistently observed across species or populations [18–20]. Because local populations can differ in genetic diversity and population structure, the genetic differences between siblings and non-siblings may also vary, potentially affecting the expression of kin discrimination [2]. Furthermore, social association preferences may be influenced by environmental conditions, including differences in the chemical cues available during rearing [21]. Thus, comparing populations which differ in genetic background under different rearing conditions may help clarify how population genetic structure and environmental context relate to kin discrimination. Additionally, kin discrimination is often discussed in the context of kin selection, but its expression alone does not establish that kin-biased associations have evolved through kin selection [2,5].

The Japanese common toad comprises two geographically differentiated lineages: the Eastern Japanese toad, *Bufo formosus*, and the Western Japanese toad, *B. japonicus* [22,23]. Historical anthropogenic introductions of *B. japonicus* into Tokyo area located eastern Japan led to hybridization with *B. formosus*, producing admixed populations [24,25]. Previous genetic analyses revealed greater genetic diversity in an admixed Tokyo population than in a homogeneous population of *B. formosus* from Nikko ([24] ; microsatellite allele richness: 2.37 and 1.50, respectively). Although kin-biased association has been reported in several anuran species, including closely related toads (e.g., in *B. americanus* [26,27]; in *B. bufo* [28]; in *B. melanostictus* [29,30]), Hase (2026) [20] found no evidence of kin recognition in *B. formosus* tadpoles from the Nikko population. However, whether kin discrimination differs between homogeneous and admixed populations, and whether such differences depend on rearing conditions, remains unclear. Hybridisation may have increased genetic differences between siblings and non-siblings within the Tokyo population, potentially facilitating kin discrimination as observed in related species.

Here, I investigated kin discrimination in tadpoles from two local populations: a homogeneous population (HomPop) of *B. formosus* from Nikko and an admixed population (HetPop) containing Eastern and Western lineages from Tokyo. I conducted two binary-choice tests under different social-rearing and water conditions. The inter-sibling test assessed kin discrimination between unfamiliar siblings and non-siblings, whereas the intra-sibling test examined familiarity-based association preferences. By comparing these responses between populations and rearing conditions, I examined whether the expression of kin discrimination varies with population genetic structure and environmental context. Understanding the conditions under which kin discrimination is expressed is an important step toward understanding the role of kin selection in the evolution of social behaviour.

## 1. METHODS

### (a) Animals and rearing

Two reproductive pairs of *B. formosus* were collected from each population: HomPop in Nikko, Tochigi Prefecture, and HetPop in Tokyo, eastern Japan (Figure 1). Tadpoles from each population were reared for two weeks either singly or in groups, using dechlorinated tap water or pond water, resulting in four rearing conditions: solitary/tap, group/tap, solitary/pond, and group/pond. Detailed procedures for producing sibling groups and rearing conditions are provided in the Supplementary Material, Figures S1 and S2.

**Figure 1.**
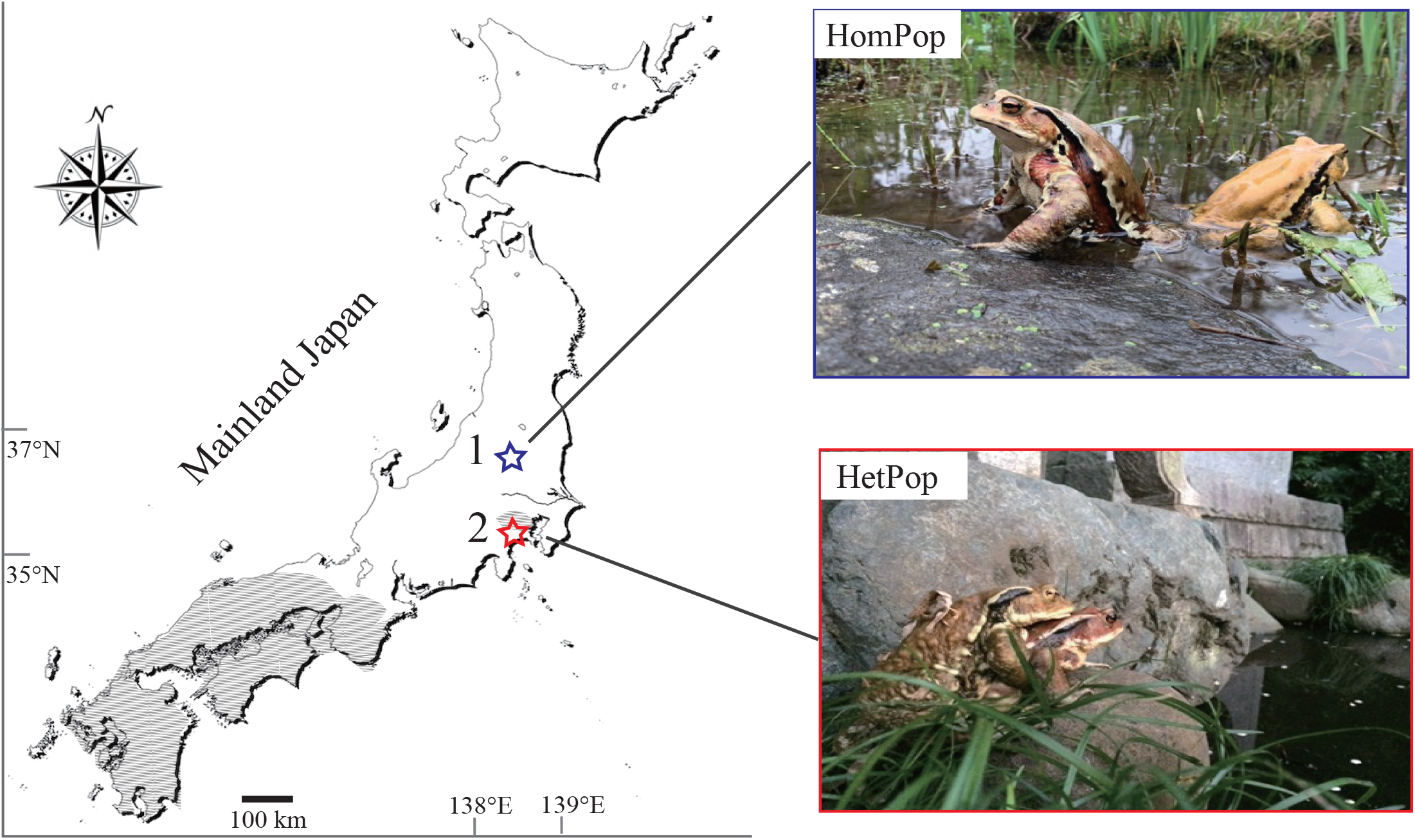
Sampling locations and breeding photographs of HomPop and HetPop.

### (b) Genotyping

To characterize the genetic backgrounds of the two populations, mtDNA (CytB) lineage and MHC class II haplotypes were determined from the four parental pairs. Genetic diversity was compared between populations. Detailed molecular procedures and sequence information are provided in the Supplementary Material, Tables S1 and S2.

### (c) Association preference tests

Association preferences were assessed using two binary-choice tests at Gosner stages 30–35 [31]. In the inter-sibling test, subjects were presented with an unfamiliar sibling and an unfamiliar non-sibling raised under the same water condition. Preference for the sibling was interpreted as phenotype-matching-based kin discrimination; comparison of group- and solitary-reared subjects allowed assessment of social learning versus self-referent phenotype matching. In the intra-sibling test, group-reared subjects were presented with a familiar sibling and an unfamiliar sibling raised under the same or different water conditions, to assess familiarity-based preferences. A total of 142 HomPop and 105 HetPop tadpoles were tested, with no subjects reused. Detailed test procedures are provided in the Supplementary Material.

### (d) Statistical analyses

Association preferences were evaluated using two-tailed paired tests. GLMMs with binomial errors were used to examine effects of population, social experience and water environment on inter-sibling preferences, and population and water conditions on intra-sibling preferences. Subject identity was included as a random effect. All statistical analyses were conducted in two steps using R version 4.3.1 [32]. Detailed model specifications are provided in the Supplementary Material.

## 2. RESULTS

### (a) Population genetic background

mtDNA analysis confirmed that HomPop comprised the Eastern lineage, whereas HetPop contained both Eastern and Western maternal lineages (Figure S3). Both populations contained three MHC haplotypes, but MHC polymorphism was higher in HetPop than in HomPop (Hd = 0.714 and 0.464, respectively; see Supplementary Material for details).

### (b) Kin discrimination and familiarity

In the inter-sibling test, association preferences differed between populations (GLMM, population effect: *P* = 0.035). Only HetPop tadpoles reared in groups in tap water significantly preferred unfamiliar siblings over unfamiliar non-siblings (*P* = 0.030; Figure 2; Table 1). No kin preference was detected in HomPop or in HetPop tadpoles reared solitarily or in pond water.

**Table 1.** Summary of the intrer-sibling test between unfamiliar siblings and unfamiliar non-siblings. Social treatment and water type are rearing conditions: reared by one (solitary) or twelve (group) per container; reared in pond water or tap water. Prefer siblings, the number of subjects who spent more on the sibling side than the non-siblings side in the choice test. P value, paired two-tailed test and bold indicates statistical significance

| Population | Social treatment | Water | Sample size | Mean time on sibling side (out of 4800s SD) | Mean time on non-sibling side (out of 4800s SD) | Prefer siblings | P value |
| --- | --- | --- | --- | --- | --- | --- | --- |
| HomPop | Group | Tap | 20 | 2357.8±628.2s | 2442.2±628.2s | 11 | 0.767 |
|  |  | Pond | 19 | 2304.5±301.7s | 2495.5±301.7s | 6 | 0.184 |
|  | Solitary | Tap | 16 | 2424.9±353.0s | 2375.1±353.0s | 7 | 0.781 |
|  |  | Pond | 16 | 2221.7±402.6s | 2578.3±402.6s | 6 | 0.097 |
| HetPop | Group | Tap | 17 | 2747.5±602.2s | 2052.5±602.2s | 11 | **0.030** |
|  |  | Pond | 11 | 2545.9±909.7s | 2254.1±909.7s | 7 | 0.606 |
|  | Solitary | Tap | 11 | 2372.5±628.3s | 2427.5±628.3s | 6 | 0.887 |
|  |  | Pond | 10 | 2356.8±491.8s | 2443.2±491.8s | 4 | 0.993 |

**Figure 2.**
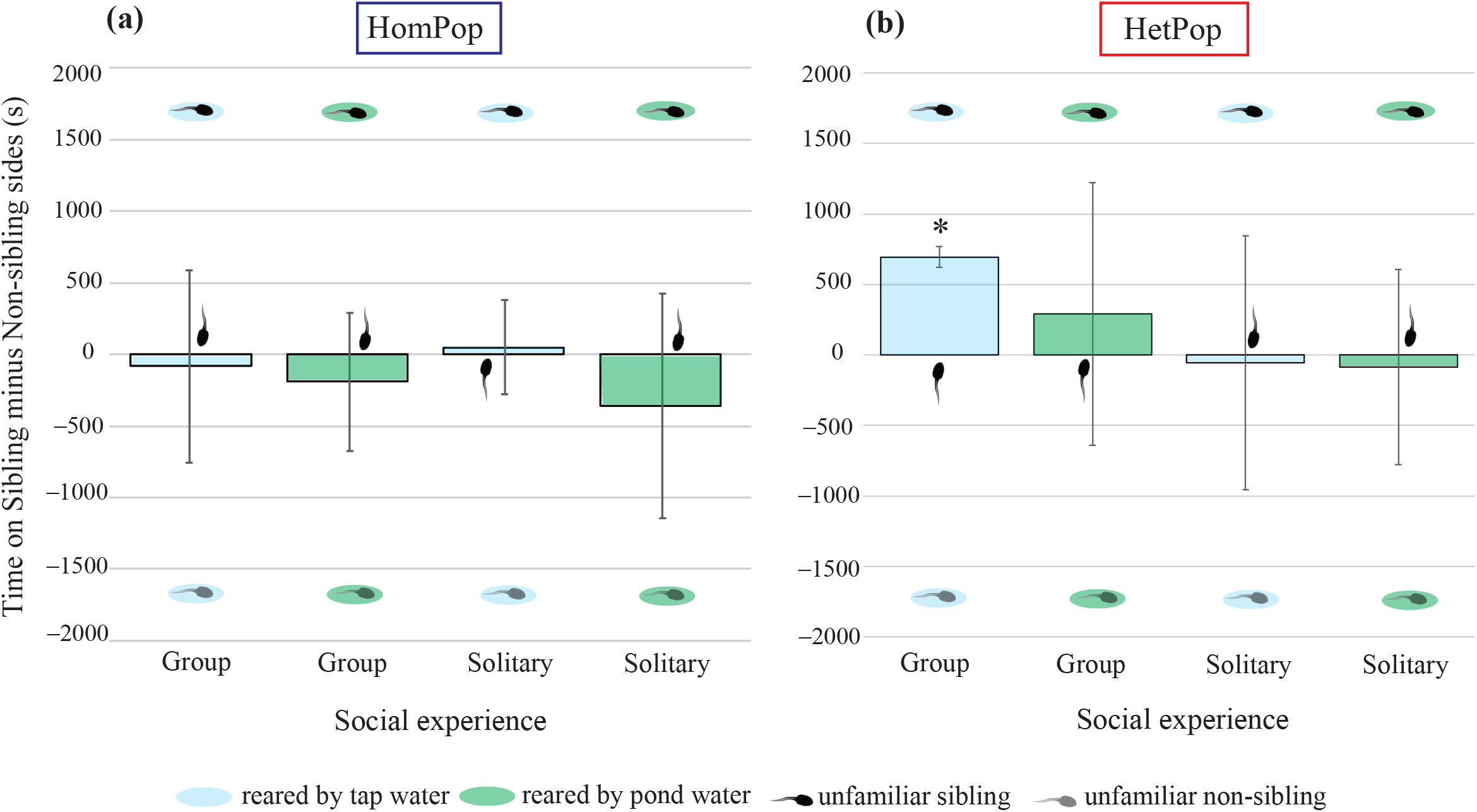
Time allocation for subject tadpoles of the inter-sibling test (mean ± 95% confidence interval) against unfamiliar non-siblings on HomPop (**a**) and on HetPop (**b**) The social treatment represents the rearing conditioning of subjects, i.e., the difference in the number of individuals in raised containers: group, twelve tadpoles; solitary, one tadpole. \**P* < 0.05 (paired two-tailed t-test)

In the intra-sibling test, HomPop tadpoles showed no familiarity preference. In contrast, HetPop tadpoles reared in tap water preferred familiar siblings when the unfamiliar sibling had been reared in pond water (P = 0.036; Fig. 3; Table 2). No explanatory variable was significant in the corresponding GLMM (details are shown in Supplementary Material).

**Table 2.** Summary of the intra-sibling test between familiar siblings and unfamiliar siblings. Water condition, the rearing water type of familiar and unfamiliar stimuli was the same (tap and tap) or the different (tap and water). Subject’s water, the water type of the subjects. Prefer familiars, the number of subjects who spent more time on the familiar sibling side than the unfamiliar sibling side in the test. P value, paired two-tailed test and bold indicates statistical significance

| Population | Water condition | Subject's water | Sample size | Mean time on familiar side (out of | Mean time on unfamiliar side | Prefer familiars | P value |
| --- | --- | --- | --- | --- | --- | --- | --- |

|  |  |  |  | 4800s SD) | (out of 4800s SD) |  |  |
| --- | --- | --- | --- | --- | --- | --- | --- |
| HomPop | Same | Tap | 18 | 2357.4±308.2s | 2442.6±308.2s | 8 | 0.565 |
|  |  | Pond | 18 | 2229.8±476.2s | 2570.2±476.2s | 6 | 0.148 |
|  | Different | Tap | 17 | 2434.2±372.0s | 2363.1±376.3s | 11 | 0.700 |
|  |  | Pond | 18 | 2450.6±273.2s | 2349.4±273.2s | 10 | 0.443 |
| HetPop | Same | Tap | 22 | 2476.0±397.1s | 2324.0±397.1s | 13 | 0.370 |
|  |  | Pond | 12 | 2451.2±783.4s | 2314.3±719.4s | 6 | 0.758 |
|  | Different | Tap | 13 | 2678.2±424.3s | 2121.2±424.3s | 9 | **0.036** |
|  |  | Pond | 9 | 2496.6±605.7s | 2303.4±605.7s | 4 | 0.645 |

**Figure 3.**
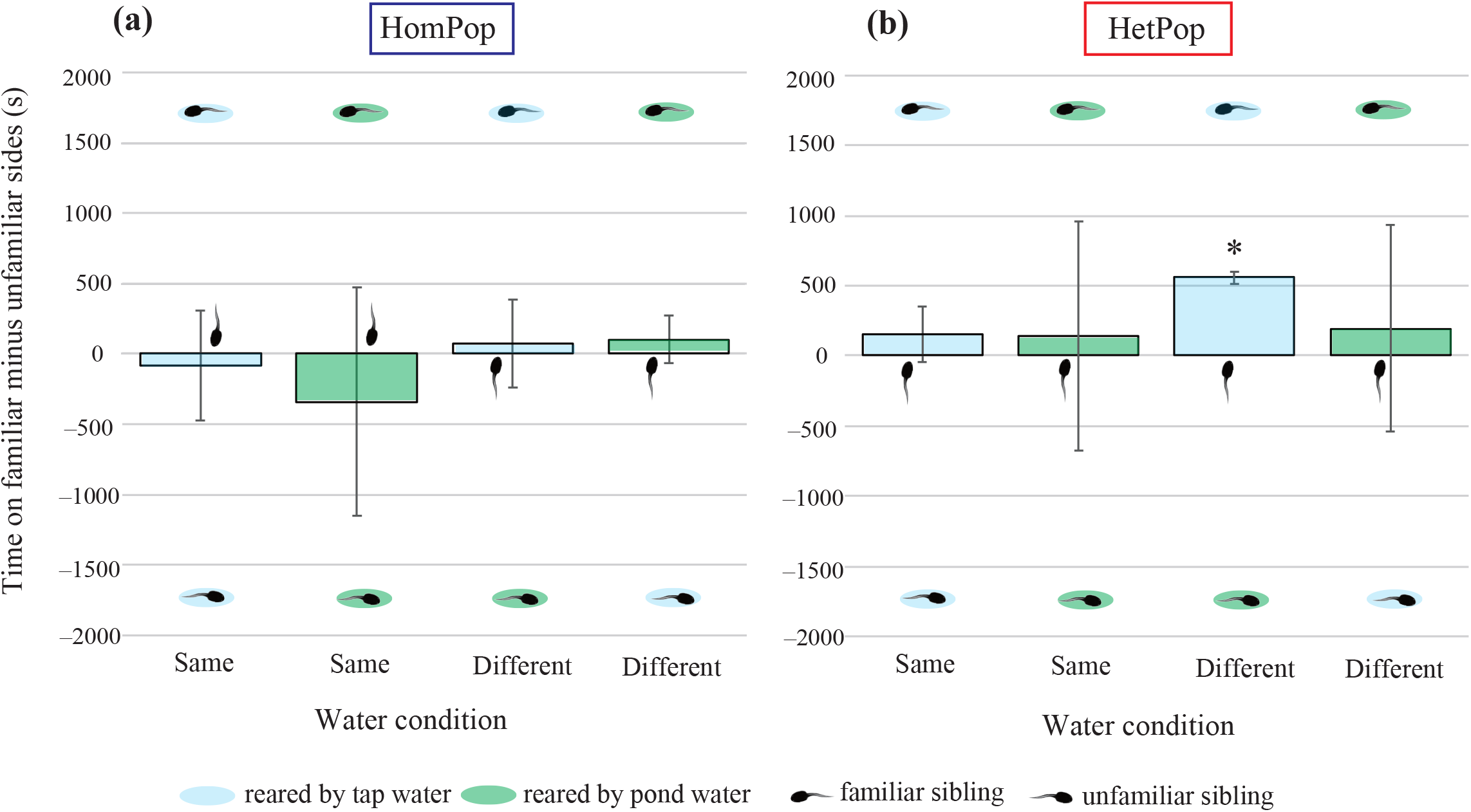
Time allocation for subject tadpoles of the intra-sibling test (mean ± 95% confidence interval) against unfamiliar siblings on HomPop (**a**) and HetPop (**b**). Water environment means the type of water in which the stimulus individuals were reared: the same, both side stimuli were raised by tap or pond water; the different, one was raised by tap water and the other by pond water. \**P* < 0.05 (paired two-tailed t-test)

## 3. DISCUSSION

Previous studies have shown that kin discrimination in anuran tadpoles varies with species, developmental experience, and environmental context [3,4,8,21,33]. Our findings extend this work by demonstrating contrasting patterns of kin discrimination between a homogeneous natural population and an admixed population of Japanese toads. Kin discrimination was detected only in group-reared tadpoles from the admixed population (HetPop) maintained in tap water, whereas it was not detected in the natural population (HomPop) under any tested condition (Figure 2). This result is consistent with Hase (2026) [20], which also found no evidence of kin recognition in tadpoles from the same natural population. The detection of kin discrimination only in group-reared HetPop tadpoles maintained in tap water suggests that its expression depends on the combination of population background, social experience, and rearing environment.

Kin discrimination in HetPop was detected only under tap-water rearing, suggesting that its expression may depend on environmental context. In the intra-sibling test, tap-water-reared HetPop tadpoles preferred familiar siblings over unfamiliar siblings reared in pond water. Although this result reflects familiarity-based association rather than kin discrimination, it suggests that differences in rearing environment may also affect social recognition. Such differences could arise from changes in the chemical cues associated with tadpoles, potentially mediated by microbial communities or other environmental factors. Together, these findings raise the possibility that population background and aquatic environment both contribute to variation in social discrimination, although the mechanisms underlying these patterns remain unresolved.

The present study suggests that kin discrimination in toad tadpoles varies with population genetic background and rearing environment rather than being a uniform characteristic of a species. Further studies examining the genetic mechanisms involved, including the potential contribution of MHC genes, and directly testing recognition cues are needed to determine whether genetic admixture contributes to variation in kin discrimination and how environmental conditions influence its expression.

### Ethics

All tadpoles were handled with care to minimise stress. The number of samples and trials was kept as low as possible. These procedures were reviewed and approved by the Ethics Committee for Animal Research of the Graduate University for Advanced Studies (no. SKD2018AR002).

## Supporting information

Supplemental methods and results, tables, figures

Raw data of association tests_ESM2

## Data accessibility

Supplemental methods and results, tables, figures, and experimental data are available online. Sequence information is deposited in GenBank (accession numbers: PQ819500– PQ819506).

## Author Contributions

KH designed the research, conducted the experiments, and prepared the manuscript.

## Acknowledgements

I am grateful to the Nikko Botanical Garden, Graduate School of Science, University of Tokyo, for permitting fieldwork. I sincerely thank Prof. N. Kutsukake for supporting the behavioural experiments at SOKENDAI, and I acknowledge Dai-chan for their advice on the kin recognition system in Bufonidae. Lastly, I express my gratitude to H. Hase for keeping the HetPop toads.

