## Supplemental methods and results, tables, figures for "Population structure and aquatic environment shape kin discrimination in toad tadpoles"

---

### 1. SUPPLEMENTARY METHODS

#### (a) Animals

Two reproductive pairs of *B. formosus* (two females and two males) were captured from breeding ponds located in HomPop in Nikko, Tochigi Prefecture, and HetPop in Tokyo, eastern Japan (Figure 1a). The collections took place in April 2017 for HetPop and March 2019 for HomPop. From these pairs, three to four sibling groups were produced (see supplementary method, Figure S1). Once the fertilised eggs had hatched, larvae were randomly selected from each egg string and gently transferred to rearing containers for the association preference test.

To investigate how social experience and the water environment affect social discrimination, the tadpoles were reared under specific conditions. For the social experience treatment, siblings were placed either singly in a 9 × 9 × 45 mm container with 50 mL of water (N = 1), or in groups of 12 in a 180 × 130 × 45 mm container with 500 mL of water (N = 12). For the water treatment, dechlorinated tap water and pond water on the university campus were prepared. Pond water was used to evaluate the impact of bacterial diversity on kin recognition, specifically through MHC-linked olfactory cues. Tadpoles were raised for two weeks in one of four conditioning environments: solitary with tap water, group with tap water, solitary with pond water, or group with pond water.

#### (b) Preparation of Sibling Tadpoles

To produce four sibling groups from two reproductive pairs, two water-filled plastic tanks (diameter 110 cm, depth 30 cm) were prepared: one tank housed one amplexus pair, and the other housed the second pair. When the female in the first tank began laying an egg string, a 15–30 cm segment of the egg string was cut off prior to the onset of male ejaculation. This egg-string segment was then placed inside a household bag-like polyethylene net (string diameter: 0.1 mm; mesh size: 13–14 threads) and transferred to the other pair's tank. Typically, toad males release sperm midway or toward the end of spawning. Once the female had finished laying her egg string and the amplexus male had released sperm, the male dismounted from the female. At this point, all toads were removed. Egg strings were left in their respective tanks for 24 hours to ensure complete fertilization—this included both the exposed (naked) eggs laid by the female and the eggs placed in the polyethylene net, which could be fertilized through the net's mesh (Figure S1a).

In total, four types of siblings were produced from the HomPop parents and three types from the HetPop parents. Figure S1b,c shows photographs of larvae in four different containers, each reflecting a distinct rearing environment.

**(c) Rearing conditioning**

Pond water was collected from the pond at The Graduate University for Advanced Studies (Hayama, Kanagawa Prefecture), then filtered through a polyethylene net (100 mesh) before use. To investigate the pond water microflora, the same procedure described in the previous study [1] was followed, including next-generation sequencing of 16S ribosomal RNA gene PCR amplicons (16S-rRNA gene-targeted amplicon sequencing) conducted at FASMAC Co., Ltd. (Kanagawa, Japan). Bacterial taxonomy (phylum/division) was analyzed using QIIME 2 [2], based on the representative sequences of operational taxonomic units defined by 97% sequence similarity. The results of the bacterial taxonomy analysis at the phylum level are shown in Figure S2.

Throughout the experiments, dechlorinated tap water and pond water were stored in 20 L polyethylene tanks in the laboratory. Solitary containers (n = 1 per container; 90 × 90 mm, 45 mm deep, holding 50 mL of water) and group containers (n = 12 per container; 180 × 130 mm, 45 mm deep, holding 500 mL of water) were placed in a large incubator (see Figure S1bc). For each sibling group (egg string), 2–4 group containers and 16–32 solitary containers were prepared, allowing 40–80 larvae to be reared. All containers were maintained at 18 °C with a 12:12 h light:dark cycle. Water was changed every two days, and tadpoles were fed daily with vegetable pellets (PLECO; Kyorin, Hyogo, Japan) until they were used for the association preference test.

**(d) Association Preference Test**

Social discrimination was evaluated adapt for an association preference test, consisting of two binary-choice tests with 2 × 2 factorial designs using four rearing conditions: the inter-sibling test (to assess kin discrimination) and the intra-sibling test (to evaluate the familiarity effect). Tadpoles at the hind limb development stage (Gosner's stages 30–35, [57] ) were used. In these binary-choice tests, *familiar* referred to siblings raised in the same container, while *unfamiliar* referred to either siblings or non-siblings raised in separate containers (raised in solitary containers, thereby unfamiliar with each other). The stimuli and subjects were paired as follows. In the inter-sibling test, to assess the effect of social experience on kin recognition, i.e., learning effect, subjects were selected from solitary or group containers, and the stimuli consisted of an unfamiliar sibling and an unfamiliar non-sibling raised under the same water condition as the subject. This design estimates the perception process of the kin recognition system: if only subjects from group containers show kin discrimination, simple phenotype-matching is indicated, whereas discrimination by subjects from solitary containers would suggest self-referent phenotype-matching. In the intra-sibling test, all subjects came from group containers to investigate familiarity effects. One stimulus was a familiar sibling, while the other was an unfamiliar sibling raised in either the same or different water conditions as the subject. A total of 142 subject tadpoles from four sibling types produced by HomPop parents, and 105 subject tadpoles from two sibling types produced by HetPop parents, were used in the binary-choice tests (no subjects were reused). Raw data from the tests have been deposited online.

All choice tests were conducted in a polypropylene tank (260 × 90 mm, 45 mm deep), partitioned into three areas: the subject arena (170 × 90 mm, 45 mm deep) and two stimulus zones on either side (45 × 90 mm, 45 mm deep). To preserve both visual (sight) and chemical (olfactory) cues, subject and stimulus tadpoles were separated by polyethylene nets (strings 0.1 mm in diameter; mesh size: 13–14 threads/$cm^2$). For each test, the tank was filled with 0.5 mL of fresh tap water. To prevent unequal microbial distribution and the accumulation of algae or bacteria on the nets, pond water was not used. At the start of each trial, the subject tadpole was gently placed in the centre of the subject arena with a 5-minute acclimation period. The subject's movement was then recorded for 40 minutes using a Panasonic HCV720-M camera, after which the sides of the stimuli were switched to eliminate positional bias, and a second 40-minute recording was made (for a total of 80 minutes per trial). The tadpoles' movement was analysed using a tracking system, UMATracker [58]. Following a previous study, the position of the subject tadpole was recorded every second by splitting the arena at the centre and tracking its two-dimensional coordinates. Association preference was calculated by comparing the total time spent on each stimulus side.

**(e) Genotyping for Parents**

Total DNA was extracted from each parent using tissue samples (approximately 3 mm toe tips) and the DNeasy Blood & Tissue Kit (Qiagen, Inc., Valencia, CA, USA) according to the manufacturer's protocol. The target mtDNA fragment (cytochrome b, 0.8 kb) was amplified by polymerase chain reaction (PCR) as described by [3,4], using previously developed primers cytbF1/Bufo (5′-ATCTGCCGAGATGTAAACAACGG-3′) and cytb177731R (5′-

TCTGYTRAGYTGGGYWAGTTTGTTCTC-3′). After alignment, the sequences were used to construct a Maximum Likelihood (ML) tree in MEGA 5.2 [5], using the GTR + Γ substitution model selected by the Akaike Information Criterion (AIC).

Partial MHC class II exon sequences (0.2 kb) were amplified by PCR using two locus-specific primers from previous studies [4,5]: 2F347 (5′-GTGACCCTCTGCTCTCCATT-3′) and R_BjMHCII (5′-CCATAGTTGTRTTTACAGWATSTCTCC-3′). After PCR, the amplified MHC class II alleles were sequenced using an ABI PRISM 310 genetic analyzer (Thermo Fisher Scientific, Inc.). In the case of heterozygous individuals, TA cloning was performed using the pGEM-T Vector System (Promega, Madison, WI, USA) and Ligation high (TOYOBO Co., Ltd., Japan); transformation was carried out with Competent high DH5 (TOYOBO Co., Ltd., Japan). After sequence alignment, six alleles were obtained from the HomPop and HetPop parents. Haplotype networks were constructed based on the number of amino acid substitutions per site among these sequences. The results for HomPop and HetPop are shown in Tables S1 and S2.

#### (d) Data Analyses

Two-tailed paired t-tests were applied to assess association preferences for siblings (inter-sibling test) or familiar individuals (intra-sibling test) by comparing the time spent in proximity to each stimulus. A significant p-value ($< 0.05$) indicated kin discrimination in the inter-sibling test, whereas in the intra-sibling test, it signified familiarity discrimination. To examine the effects of rearing conditions and population genetic structure on association preferences in both the inter- and intra-sibling tests, general linear mixed models (GLMMs) with binomial error distributions were used. In the inter-sibling test, the proportion of time spent near the sibling stimulus (out of a total of 4,800 seconds) served as the response variable, and subject identity was treated as a random factor. Fixed explanatory variables included population (HomPop or HetPop), social experience (solitary or group), water environment (tap or pond), and the two-way interaction of social experience and water environment. In the intra-sibling test, the proportion of time spent near the familiar-sibling stimulus (out of a total of 4,800 seconds) was the response variable, and subject identity was again treated as a random factor. Fixed explanatory variables included population, the water type of familiar siblings (tap or pond), the water type of unfamiliar siblings (same or different), and the two-way interaction between these water types.

### 2. SUPPLEMENTARY RESULTS

**(a) Genotyping on Parents**

Figure 1b shows the phylogenetic relationship of captured toads from two populations based on mtDNA (CytB, 0.8 kb). All toads from HomPop belonged to the Eastern clade, whereas the toads from HetPop were divided into two clades: two males and one female were in the Western clade, while the other female was in the Eastern clade. This result is consistent with previous studies [42,43], indicating that HomPop is a naturally distributed population of *formosus*, whereas HetPop is a heterogeneous local population containing both native *formosus* and introduced *japonicus* maternal lineages. Figure 1c depicts the haplotype network of amino acid sequences of MHC alleles identified from parents in HomPop and HetPop. Both populations contained three MHC haplotypes, but their diversity (haplotype and nucleotide polymorphism) was higher in HetPop than in HomPop. The calculated values for HomPop and HetPop were as follows: Hd (mean ± SD) = 0.464 ± 0.20 and Hd = 0.714 ± 0.12, k = 5.964 and k = 8.714, and NonSyn/Syn ratios of 0.807 and 2.843, respectively. Sequence information is deposited in GenBank (accession numbers: PQ819500– PQ819506).

**(b) Preference Tests in Tadpoles**.

Table 1 summarises the inter-sibling test. Each population was examined to spend time on the unfamiliar sibling and non-sibling sides under different social and water environment conditions (Figure 2ab). The association preferences for the siblings of HomPop and HetPop displayed a marked difference. GLMM analysis detected a significant effect solely for the population factor ($z = -2.110$, $P = 0.035$). In contrast, no effects were observed for social experience ($z = -0.669$, $P = 0.504$), water environment ($z = 0.970$, $P = 0.332$), or their two-way interaction ($z = -0.145$, $P = 0.885$). Consequently, only tadpoles originating from HetPop parents, raised in groups and tap water, showed a significant preference for associating with unfamiliar siblings over non-siblings, suggesting phenotype-matching-based kin recognition ($P = 0.03$; Table 1, Figure 2b). Tadpoles from the naturally occurring HomPop exhibited no kin preference for unfamiliar siblings; no evidence for either self-referent or phenotype-matching-based kin recognition was found (Figure 2a).

Table 2 summarises the intra-sibling test. Subject tadpoles were evaluated in four designs for each population, reflecting differences in the water environment between the subject and the unfamiliar stimulus (Figure 3). Tadpoles from HomPop, which showed no preference in the inter-sibling test, likewise exhibited no significant preference for familiar siblings over unfamiliar ones, irrespective of the water environment (Figure 3a). By contrast, HetPop tadpoles raised in tap water significantly preferred familiar siblings when unfamiliar ones had been raised in pond water (Figure 3b). Unlike in the inter-sibling test, none of the explanatory variables showed significant effects in the GLMM analysis (Population, z = -1.946, $P$ = 0.052; water condition of familiar ones, z = 0.308, $P$ = 0.758; water condition of unfamiliar ones, z = 1.069, $P$ = 0.285; two-way interaction, z = 0.295, $P$ = 0.768).

### 3. SUPPLEMENTARY TABLES AND FIGURES

**Table S1**. Estimates of evolutionary divergence between sequences of MHC class II PBR alleles from HomPop parents. Standard error estimate(s) are shown above the diagonal. Analyses were conducted using the Poisson correction model [6]. The rate variation among sites was modeled with a gamma distribution (shape parameter = 1). The analysis involved 3 amino acid sequences. All positions containing gaps and missing data were eliminated. There were a total of 63 positions in the final dataset. Evolutionary analyses were conducted in MEGA [4]

| **HomPop** | Allele 1a | Allele 1b | Allele 1c |
|---|---|---|---|
| Allele 1a | | 0.045 | 0.067 |
| Allele 1b | 0.105 | | 0.067 |
| Allele 1c | 0.189 | 0.189 | |

**Table S2.** Estimates of evolutionary divergence between sequences of MHC class II PBR alleles from HetPop parents. Standard error estimate(s) are shown above the diagonal. Analyses were conducted using the Poisson correction model [5]. The rate variation among sites was modeled with a gamma distribution (shape parameter = 1). The analysis involved 3 amino acid sequences. All positions containing gaps and missing data were eliminated. There were a total of 63 positions in the final dataset. Evolutionary analyses were conducted in MEGA [4].

| **HetPop** | Allele 2a | Allele 2b | Allele 2c |
|---|---|---|---|
| Allele 2a | | 0.069 | 0.04 |
| Allele 2b | 0.189 | | 0.08 |
| Allele 2c | 0.086 | 0.235 | |

**Figure S1.** Photos of tadpole raising. Fertilization of naked eggs and eggs in polyethene net bag (a). Conditioning of water environment for group and solitary raising: tap water (b) and pond water (c)

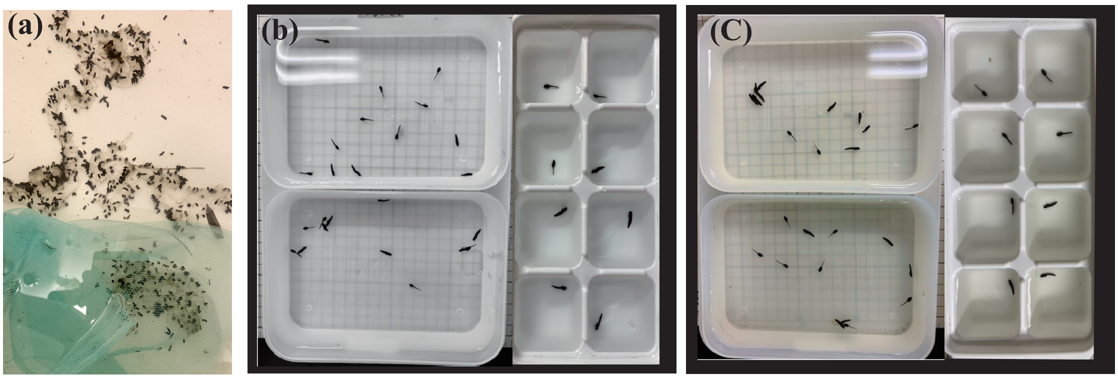


**Figure S2.** Relative abundances of bacterial taxonomy at the phylum level of the pond water sampled in 2017 and 2019

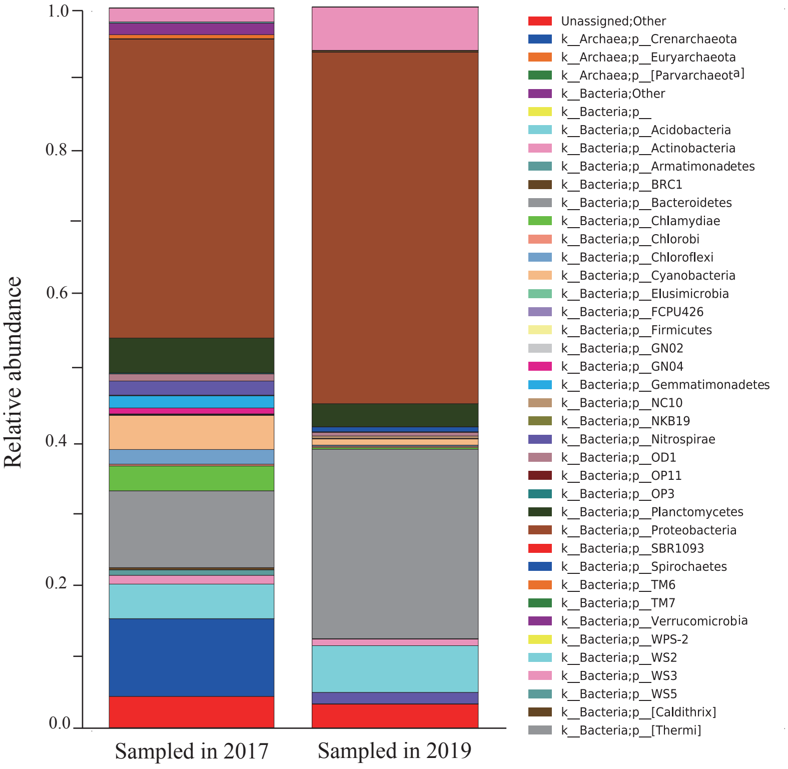

**Figure S3**. (**a**) Maximum Likelihood (ML) tree based on mtDNA Cyt*B* (0.8 kb) from all captured toads, constructed using the GTR + Γ model. Support values for internal nodes (>70) were inferred from 1,000 bootstrap replicates. Toads captured in the present study are shown in bold, while other sequences are references from previous study (accession numbers: AB597912–AB597929). Labels for Hom and Het indicate haplotypes originally derived from individuals sampled from the same HomPop and HetPop populations in earlier studies [40,41]. (**b**) Haplotype networks based on MHC class II PBR amino acid sequences obtained from the captured parents. Each circle represents an allele, with circle diameter reflecting the number of alleles (see the supporting Table S1 for calculation details). Female1.1, Female1.2, Male1.1, and Male1.2 belong to HomPop, while Female2.1, Female2.2, Male2.1, and Male2.2 are from HetPop.

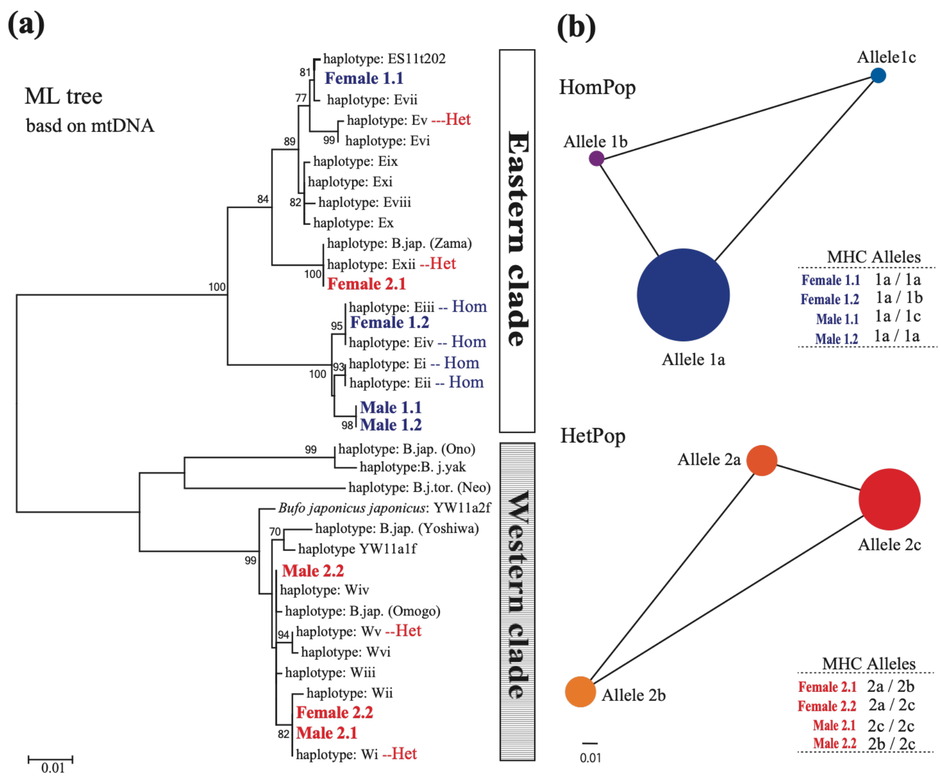
